# Temporal Dynamics of the Origin and Domestication of Sweet Potato and Implications for Dispersal to Polynesia

**DOI:** 10.1101/309799

**Authors:** Pablo Muñoz-Rodríguez, Tom Carruthers, Robert W. Scotland

## Abstract

In a study focussed on the origin of *Ipomoea batatas* (L.) Lam. (the sweet potato) and its close relatives [1], the authors claimed that a specimen of sweet potato collected by Banks and Solander in 1769 in Polynesia was distinct and along with some other varieties had levels of genetic variation of more than 100,000 years. This timeframe, along with several other lines of evidence, was interpreted, by the authors, as questioning human-mediated transport of the sweet potato to Polynesia within the last 1,000 years. This aspect of the paper received widespread media coverage as well as a number of critical remarks, and correspondence. The criticism questioned whether claims in the paper were justified. Here, three authors from the paper take the opportunity to fully explain the reasoning for calling into question the need to invoke human mediated transport of the sweet potato to Polynesia.

## Iterative nature of long-distance dispersal and storage root evolution in *Ipomoea*

Our study of sweet potato and its wild relatives is part of a much larger taxonomic [2–10] and phylogenetic study of *Ipomoea* worldwide, to be published later this year. During the last four years, we have become aware of the prevalence of naturally occurring long distance dispersal within and across the phylogenetic breadth of *Ipomoea* and we provide two examples in the paper [1]. The significance of this highly iterative process is that disjunct distribution patterns of closely related species (e.g. disjunction patterns between Polynesia and the American continent) are not uncommon in *Ipomoea*, and therefore a species native to America but also present in Polynesia (whether it is a crop or not) is not that surprising. However, the prevalence of naturally occurring long distance dispersal in *Ipomoea*, in itself, does not preclude the sweet potato arriving in Polynesia via human mediated dispersal within the last 1,000 years. What we claim, though, is that the probability of it dispersing naturally to Polynesia is already high given the prevalence of naturally occurring dispersal of close relatives across vast distances, and becomes even more so within the timeframe since it diverged from its closest relative, *I. trifida*.

In addition, during our field studies [2–10] we became aware of one other aspect of *Ipomoea* biology: the independent evolution of tubers in different species in various clades across the phylogenetic breadth of the genus. Storage roots have evolved on, at least, 10 separate occasions within *Ipomoea* where most species do not posses a tuber. This observation highlights the relative ease and prevalence of root enlargement in this group of plants. In contrast, prior domestication literature on sweet potato explicitly assumes a human-mediated process of selection for tuberous roots from non-tuberous species [see for example 11–13]. In the paper, we implicitly set out to test these two hypotheses as rigorously as possible. We wanted to explore whether our data were in support of human domestication of the sweet potato’s storage root or whether this structure is a pre-adaptation that predisposed *Ipomoea batatas* to cultivation in one or more than one biogeographical and cultural context. We attempted this within a paper that had many elements centred on a comprehensive phylogenetic study of the sweet potato and all of its close relatives and maybe our test of the timeframe for evolution of the tuber could have been more explicit, in which case the connection between the time-frame and the Polynesian part of the paper would have been more clear.

However, the relative regularity within which distantly related species of *Ipomoea* have evolved storage roots, coupled with a pre-human timeframe for the origin of *Ipomoea batatas* from its closest relative (*I. trifida*), provides a compelling challenge to the view that the sweet potato root is mostly a product of human domestication. It is likely that sweet potato’s characteristic root was already present when it was taken into cultivation with minimal subsequent phenotypic changes, a scenario that has been suggested for other crops in the Neotropics [14].

Taken together, the three pieces of evidence —the origin of *I. batatas* more than 0.8 million years ago, the iterative nature of storage root evolution and the prevalence of naturally occurring long distance dispersal within *Ipomoea*— provide a much longer and older timeframe within which to postulate sweet potato dispersal to Polynesia compared with that of domestication of sweet potato within the last 8000 years and it being taken to Polynesia by humans in the last 1000 years.

It was in this context that we decided to sequence the specimen of sweet potato collected by Joseph Banks and Daniel Solander in 1769, to investigate how the DNA sequence data from that specimen fitted the timeframe we had developed from other data.

## Banks & Solander Specimen and Possible Contamination

The fragment of leaf tissue of the Banks and Solander specimen that we obtained from the Natural History Museum in London was processed and prepared two years after the rest of our samples. The libraries were prepared in a laboratory with no history of involvement with our project and the sequencing-run contained no other *Ipomoea* samples. Given the fact that 1) the assembled chloroplast genome of this specimen clearly places it within one of the two sweet potato lineages we identified (the same group to which it was assigned in a previous study using only a small chloroplast region [15]); 2) it can be distinguished from all other samples in our study; and 3) there are other samples which seem to have accumulated a similar level of mutations throughout time (see for example Figure 7A in [1]), it means that the probability of cross-contamination from other *Ipomoea* samples is very small. We followed the standard protocol in systematics for extracting DNA from herbarium specimens (including one other *Ipomoea* specimen collected more than 100 years ago) and did not treat them as ancient DNA. In retrospect, and given the sweet potato and Polynesia is a topic that generates such strong reactions, we acknowledge that treating the specimen as an ancient DNA sample would have been prudent and preferable.

## Chloroplast and Nuclear Data from Banks and Solander

When sequencing the Banks and Solander specimen, our aim was to assemble the whole chloroplast genome and to check what chloroplast lineage it belonged to, for which we used genome skimming. Because this technique randomly retrieved fragments of the nuclear regions previously analysed in our study, we decided to try to assemble those fragments and incorporate them to our nuclear phylogenies, aiming to obtain some insights from the nuclear genome. Despite using a very conservative approach when assembling and analysing the nuclear regions (explained in [1]), we were aware that our chloroplast data had much higher read coverage and was more robust than the lower quality of data obtained for the nuclear data, and for that reason our analyses to estimate divergence times were conducted using the chloroplast data only (see below). We considered presenting the chloroplast data alone and not including the nuclear data but we felt that including all the data was both more open and transparent. In retrospect, we should have presented the chloroplast data but not the nuclear data. In particular, our supplementary Figure S7 has an inflated branch length which is most probably an artefact due to low read coverage and sequencing error. Rather than explore the nuclear data further, we focussed on the chloroplast. We fully concede that reporting this preliminary nuclear data, in the way we did, was a mistake. However, because we knew the nuclear data was incomplete we based our two claims relevant to the Banks and Solander specimen on the chloroplast. We claimed that the chloroplast genome was distinct from other samples in our study and that, given our time-calibration, the Banks and Solander specimen diverged from other samples in our study outside of a timeframe for human domestication, more than 100,000 years ago. We stand by these two claims. This does not mean the sweet potato was not transported to Polynesia by humans, but simply that the Banks and Solander specimen, along with other specimens in our study, had levels of genetic variation that placed them in a much older time-frame (Figure 1) compared with what would be expected if the tuber was a product of human domestication and if it were brought from America within the last 1,000 years.

**Figure 1.**
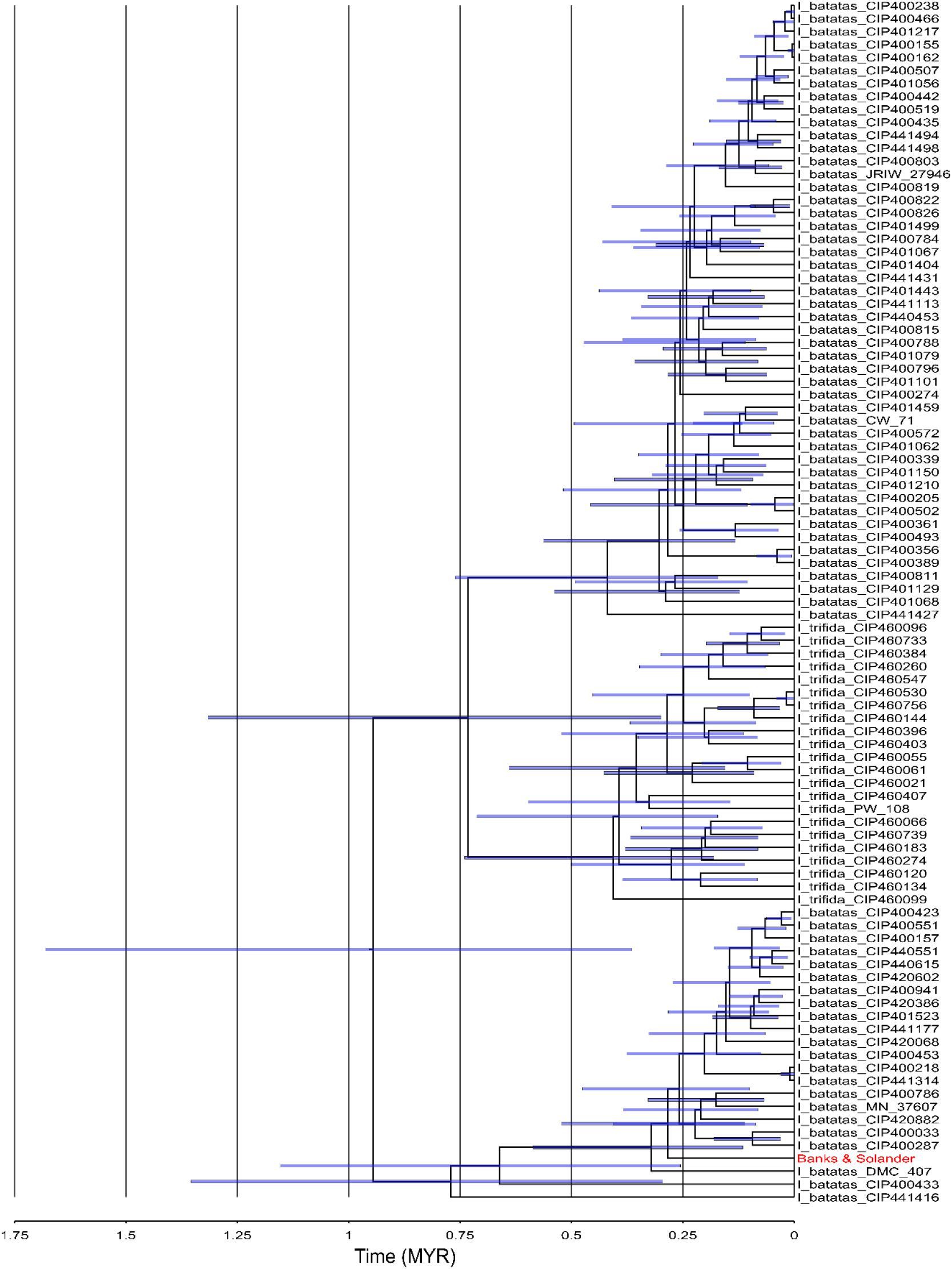
Chloroplast gene tree generated from our multispecies coalescent analysis of 94 specimens of *Ipomoea trifida* and *Ipomoea batatas*. This tree illustrates the coalescent times between individual chloroplast samples, with blue error bars representing the 95% highest posterior density. Note that coalescent times between some *Ipomoea batatas* specimens are likely to significantly predate humans, whilst coalescent times for other samples overlaps with the human era. X-axis scale in millions of years, for terminal taxa labels see [1].

## Divergence Times

With respect to divergence time estimation, our investigation was primarily concerned with estimating whether *Ipomoea batatas* evolved in pre-human times. To assess this in the most robust manner, we estimated a minimum age for the divergence between *Ipomoea batatas* and its closest relative, *Ipomoea trifida*. Secondarily, we were interested in estimating the extent to which diversity within *Ipomoea batatas* evolved in pre-human times. This analysis was based on 94 specimens of *Ipomoea batatas* and *Ipomoea trifida*, one of which was the *Ipomoea batatas* specimen collected by Banks and Solander.

In all our divergence time estimates, we used a 67.34 million year calibration for the divergence between Convolvulaceae and Solanaceae. This age is derived from an angiosperm-wide study that incorporated 136 fossil calibrations [16]. We consider this calibration an underestimation of the age of this node; the method used to infer this date is likely to bias divergence time estimates to younger ages, whilst fossil discoveries within Solanaceae imply a drastically older age for this node [17]. Together, this highlights that our divergence time estimates are biased to young ages. This provides the most robust test of whether *Ipomoea batatas*, and diversity within it, evolved long before the timeframe for human domestication.

With respect to the molecular data we used, our analysis of the entire batatas group (to demonstrate when *Ipomoea batatas* diverged from its closest relative, *Ipomoea trifida*) was conducted with 21 nuclear genes and independently with plastome data. These datasets are from entirely different genomes and thus likely exhibit different patterns of among lineage molecular evolutionary rate heterogeneity. The fact that our conclusions are broadly consistent with both datasets suggests our findings are robust to inherent problems in estimating molecular evolutionary rates [18,19].

Our results demonstrate that *Ipomoea batatas* evolved at least 800,000 years ago. Unsurprisingly, within *Ipomoea batatas*, we demonstrate that some diversity evolved hundreds of thousands of years ago, whilst some is likely to have evolved far more recently (Figure 1). With respect to the Banks and Solander specimen, it likely diverged from its closest relative over 100,000 years ago according to the chloroplast data. Nonetheless, we recovered the same general result —a significant amount of diversity within *Ipomoea batatas* evolving in pre-human times whilst some evolving far more recently— regardless of whether we included the Banks and Solander specimen in our analyses. Given the timeframe we present, and the pervasive influence of long distance dispersal on plant distributions (including two examples in close relatives of *Ipomoea batatas* presented in the paper), we consider it likely that *Ipomoea batatas* dispersed naturally across the Pacific.

Of further interest, we note that our estimate for the divergence between *Ipomoea trifida* and *Ipomoea batatas* (>800,000 years) is congruent with that obtained in a recent paper based on an *Arapidopsis* mutation rate calculated over 30 generations [20]. We did not consider it worthwhile to repeat this analysis. In our analyses, we simultaneously estimate a mutation rate when estimating divergence times. We could have constrained our model to a set value, such as that obtained by Yang *et al*., [20]. This would have resulted in more precise divergence time estimates, but there is no reason to believe these estimates would be more accurate. Further, this approach would not have addressed the primary source of error in divergence time estimates when analysing genomic scale datasets — namely among lineage (mutation) rate heterogeneity [18,19].

## In Conclusion

Our claims are that sweet potato and its storage root diverged from its sister species 800,000 years ago. This exact date, we take with a pinch of salt, as we are aware of the many issues associated with divergence time estimates. However, we deliberately biased our analyses to be as young as possible, but even so, the samples of *Ipomoea batatas* that we analysed demonstrate a time-scale that is compatible with a pre-human origin of the sweet potato root (Figure 1) and certainly much older than the timeframe for domestication scenarios over the last 8000 years. Furthermore, in a genus in which long distance dispersal is prevalent, the timescale for the sweet potato to naturally disperse to Polynesia is a very long one and this means that naturally occurring dispersal is much more likely the way that the sweet potato reached Polynesia.

